# Mnemonic Demand Reconfigures the Neural Architecture of Memory Encoding

**DOI:** 10.64898/2026.09.03.749199

**Authors:** Chong Zhao, Edward K. Vogel, Monica D. Rosenberg

**Affiliations:** Department of Psychology, University of Chicago, Chicago, Illinois 60637, USA; Institute for Mind and Biology, University of Chicago, Chicago, Illinois 60637, USA; Neuroscience Institute, University of Chicago, Chicago, Illinois 60637, USA

**Keywords:** Visual Working Memory, Visual Long-Term Memory, Neural Geometry, Interelectrode Correlation, EEG

## Abstract

Memory encoding must accommodate increasingly demanding loads, yet it remains unclear whether the underlying brain state is preserved or adaptively reconfigured as load increases. Here, we recorded EEG from 111 participants performing a visual memory task in which they encoded lists of real images ranging from 1 to 128 items, spanning low to high mnemonic demands, followed by a recognition test. We measured the similarity of set-size-specific interelectrode correlation patterns to a low-load reference. Similarity decreased monotonically as list length increased: patterns at set sizes below 4 remained close to the low-load configuration, whereas set sizes of 8 and above progressively shifted toward a distinct high-load configuration. This transition was captured by a single rotating eigenvector, revealing a low-dimensional trajectory from low- to high-load neural geometry. Smaller rotation angles from the low-load template predicted better memory performance at high loads. Together, these findings provide evidence for adaptive reconfiguration and reveal a structured, low-dimensional neural trajectory through which encoding architecture changes as mnemonic demands increase.

## Introduction

Memory encoding is often treated as an operation applied to individual events: incoming information is selected, represented, and either succeeds or fails to leave a durable trace. Yet the operation performed on an event may depend critically on the state of the memory system when that event arrives. The first object in an episode is encountered when little else must be remembered; the same object encountered after dozens of preceding events arrives in a system carrying a substantial mnemonic history. This raises a basic question that is difficult to address when working memory (WM) and long-term memory (LTM) are studied separately: as mnemonic demand accumulates, does the brain continue to encode new events using a stable neural state whose effectiveness varies, or does the organization of encoding itself change?

The enormous difference in capacity between WM and LTM provides a natural way to expose this question. Visual WM can actively maintain only a small number of objects, commonly estimated at approximately three to four items ^1,2^, with neural measures of WM load showing similar capacity limits ^3^. Visual LTM, in contrast, can retain thousands of distinct objects with substantial detail ^4–6^. Thus, as a sequence extends from one item to dozens or hundreds, the mnemonic circumstances under which each new event is encoded change dramatically. Early information can coexist within an actively maintained set, whereas successful retention of an extended sequence necessarily requires information to survive beyond that limited active state. This distinction does not require WM and LTM to be isolated stores, nor does it imply that any particular list length engages one system exclusively. Rather, the capacity limit provides a principled point at which the demands placed on encoding change.

One possibility is that increasing demands reduce the efficacy of a common encoding state without fundamentally changing its organization. According to a **repeated-loading account**, the same core operations continue to select, represent, and establish durable traces for successive items, while the effectiveness of those operations changes with recent mnemonic history. This view is compatible with activation-based models ^7^, in which memory performance varies continuously with representational strength, activation, interference, and contextual binding rather than through transitions between discrete encoding systems. Similarly, the Source of Activation Confusion framework and its resource-limited extensions propose that episodic encoding depends on binding information to context, with the resources supporting those bindings shaped by prior processing demands and recovery over time ^8–10^. Thus, repeated loading can progressively weaken encoding across an episode without requiring a fundamental change in the underlying neural coordination architecture.

A different possibility is that increasing mnemonic demand triggers an **adaptive reconfiguration** of the encoding state itself. Once active maintenance can no longer accommodate the accumulating episode, the relative contributions of selection, maintenance, contextual updating, and durable storage may change, producing a new organization of large-scale neural coordination. Such a transition need not constitute an abrupt switch between separate WM and LTM systems. Instead, encoding may follow a graded trajectory in which its neural state progressively departs from the low-load configuration as mnemonic history accumulates. Critically, this account predicts a systematic change in the relational structure of neural activity rather than a weaker or noisier expression of the same brain state: higher-demand states should not be fully recoverable by simply scaling, attenuating, or adding noise to the low-load architecture.

In the current study, we recorded EEG from 111 participants performing a visual memory encoding task with set sizes ranging from 1 to 128 items (**Fig. 1**). We characterized large-scale neural organization using interelectrode correlation (IC) patterns, which capture the coordinated structure of activity across the scalp ^11–13^. We tested two accounts of encoding under increasing mnemonic demands: **repeated loading**, which predicts that the same neural architecture is progressively loaded as set size increases, and **adaptive reconfiguration**, which predicts that the neural pattern associated with encoding is reorganized as demands exceed WM capacity. We therefore asked whether increasing mnemonic demand leaves the underlying IC architecture unchanged, as predicted by repeated loading, or alters its organization, as predicted by adaptive reconfiguration. We further tested whether these changes track memory load and predict individual differences in memory performance.

**Fig 1.**
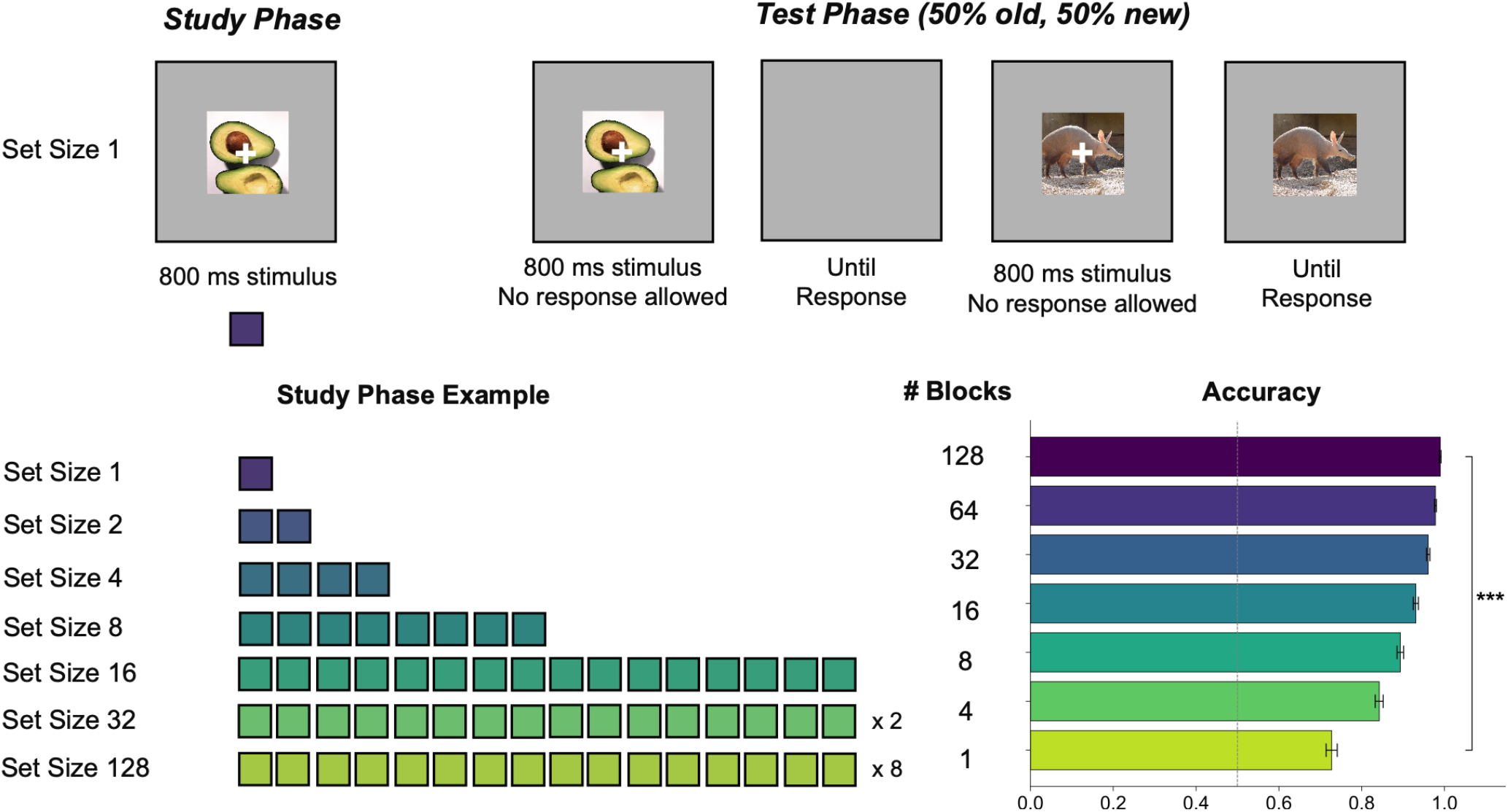
Visual Recognition Memory Paradigm with set size 1, 2, 4, 8, 16, 32 and 128 with EEG.

**Fig 2.**
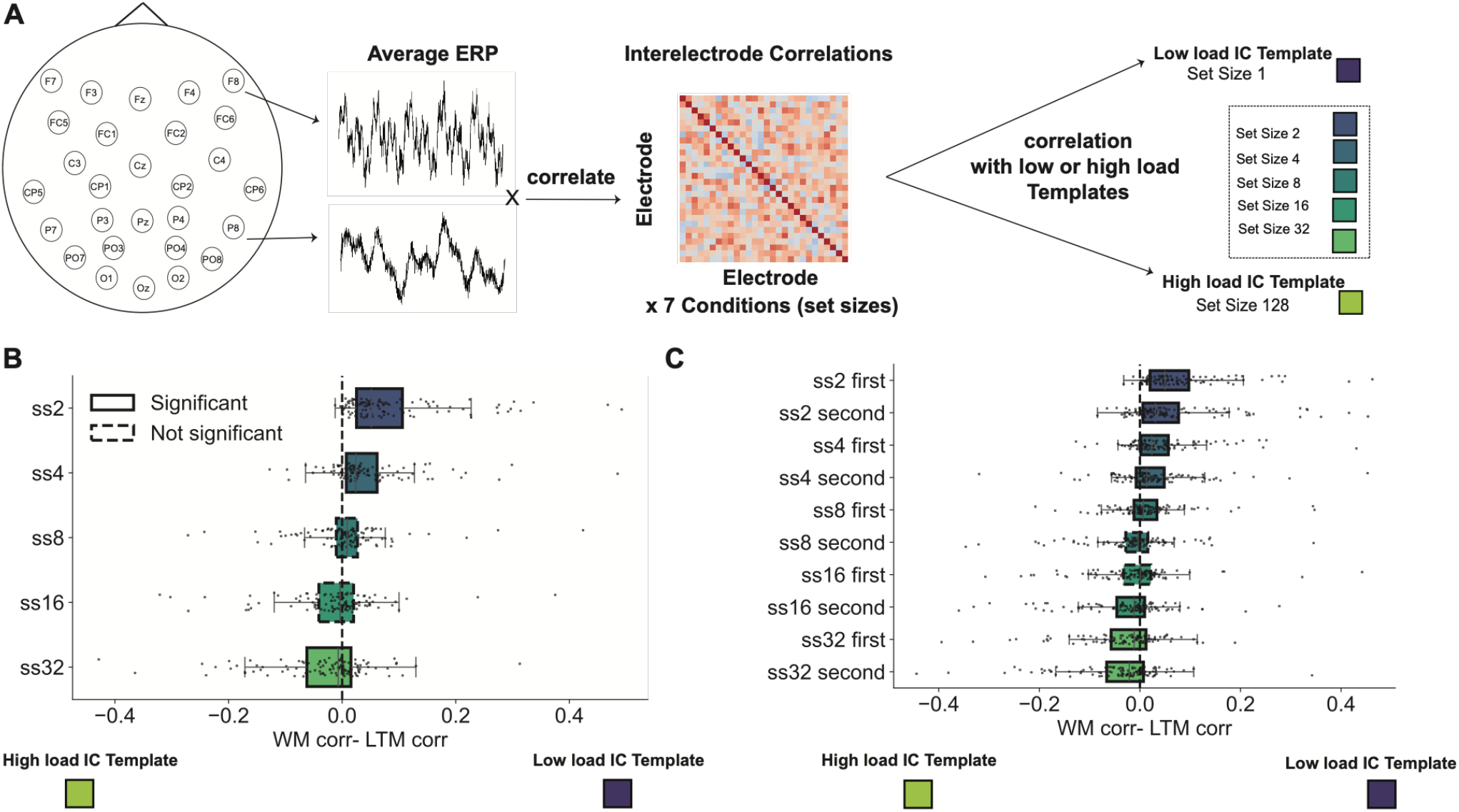
Interelectrode Correlation (IC) similarities revealed a shift from low-load to high-load to pattern as set size increases.

## Results

### Interelectrode correlation patterns track adaptive reconfiguration as mnenomic demands increase

111 participants completed 128 encoding trials at each of seven set sizes (1, 2, 4, 8, 16, 32, and 128 images), yielding 896 encoding trials per participant. By systematically increasing list length from within WM capacity to far beyond it, we tested two accounts of encoding under increasing mnemonic demands: **repeated loading**, which predicts that the same encoding architecture is progressively loaded as list length increases, and **adaptive reconfiguration,** which predicts a shift to a distinct encoding architecture as demands exceed WM capacity.

To characterize neural encoding, we used interelectrode correlation (IC) patterns, computed as cross-correlations across scalp electrodes from trial-averaged EEG time series ^11–13^. For each participant, we dervied an IC template for each set size. The set size 1 template provided a baseline for *low-load* encoding, whereas the final 64 trials of the set size 128 condition provided a *high-load* template that minimized contamination from early-list processing.

The two accounts make distinct predictions. **Repeated loading** predicts that longer lists are encoded by repeatedly applying the same architecture used for shorter lists. Thus, increasing set size should primarily increase the amount of loading within a stable encoding configuration, with IC patterns remaining relatively similar across set sizes (for example, a 16-item list should resemble repeated applications of the encoding configuration observed for a 4-item list). **Adaptive reconfiguration**, in contrast, predicts that accumulating mnemonic demands eventually alter the organization of the encoding state itself, changing IC-pattern geometry as list length increases beyond WM capacity. We therefore quantified the similarity of each set-size pattern to the low- and high-load templates.

We found evidence consistent with **adaptive reconfiguration.** At set sizes 2 (t(110) = 9.57, p < 0.001) and 4 (t(110) = 5.89, p < 0.001), neural patterns were more similar to the low-load template. At set sizes 8 (t(110) = 1.01, p = 0.31) and 16 (t(110) = −1.73, p = 0.09), patterns became intermediate between the low- and high-load templates. By set size 32, patterns were significantly more similar to the high-load template (t(110) = −3.40, p < 0.001). Thus, rather than remaining stable across set sizes as predicted by repeated loading, neural representations progressively shifted toward a distinct high-load configuration.

Because longer lists necessarily contain early items processed under lower mnemonic demands, we next asked how this transition unfolded within lists. To determine whether the encoding state changed as mnemonic history accumulated while keeping the amount of neural data used for each estimate constant, we divided each condition into early and late halves. This resulted in 64 encoded items contributing to each estimate at every set size. At set sizes 2 and 4, both halves remained more similar to the low-load template (t(110)s > 6.70, ps < 0.001), consistent with a stable encoding architecture within WM capacity. At set size 8, the first half remained more similar to the low-load template (t(110) = 2.08, p = 0.04), whereas the second half showed an intermediate pattern (t(110) = −1.56, p = 0.12), indicating the onset of reconfiguration after WM capacity was reached. At set sizes 16 and 32, this shift became progressively stronger: later items showed greater similarity to the high-load template (t(110) = −3.29, p = 0.001 and t(110) = −4.54, p < 0.001, respectively), whereas earlier items retained mixed or low-load patterns.

Together, these findings indicate that increasing mnemonic demands do not simply load a fixed encoding architecture. Instead, as demands accumulate beyond WM capacity, neural representations progressively reorganize toward a distinct encoding configuration.

### Adaptive Reconfiguration Is Captured by a Low-Dimensional Neural Transformation

A remaining question is how large-scale neural coordination shifts across memory encoding modes. The interelectrode correlation (IC) space underlying this transition is high- or low-dimensional. If the low-to-high load shift is encoded in a high-dimensional IC space, then no single eigenvector would suffice to capture it. The transition would instead be distributed across many orthogonal dimensions. Alternatively, if the first eigenvector alone accounts for the systematic change in IC patterns across list lengths, this would imply that the reconfiguration follows a constrained trajectory through neural state space.

To distinguish between these possibilities, we applied Singular Value Decomposition (SVD) to interelectrode correlation matrices computed separately for each of the seven set-size conditions (set sizes 1, 2, 4, 8, 16, 32, and 128 items; **Fig. 3A**). SVD decomposes the full pattern of pairwise electrode correlations into orthogonal components ranked by their explained variance, with the first eigenvector capturing the dominant axis of coordinated neural activity across recording sites.

**Figure 3.**
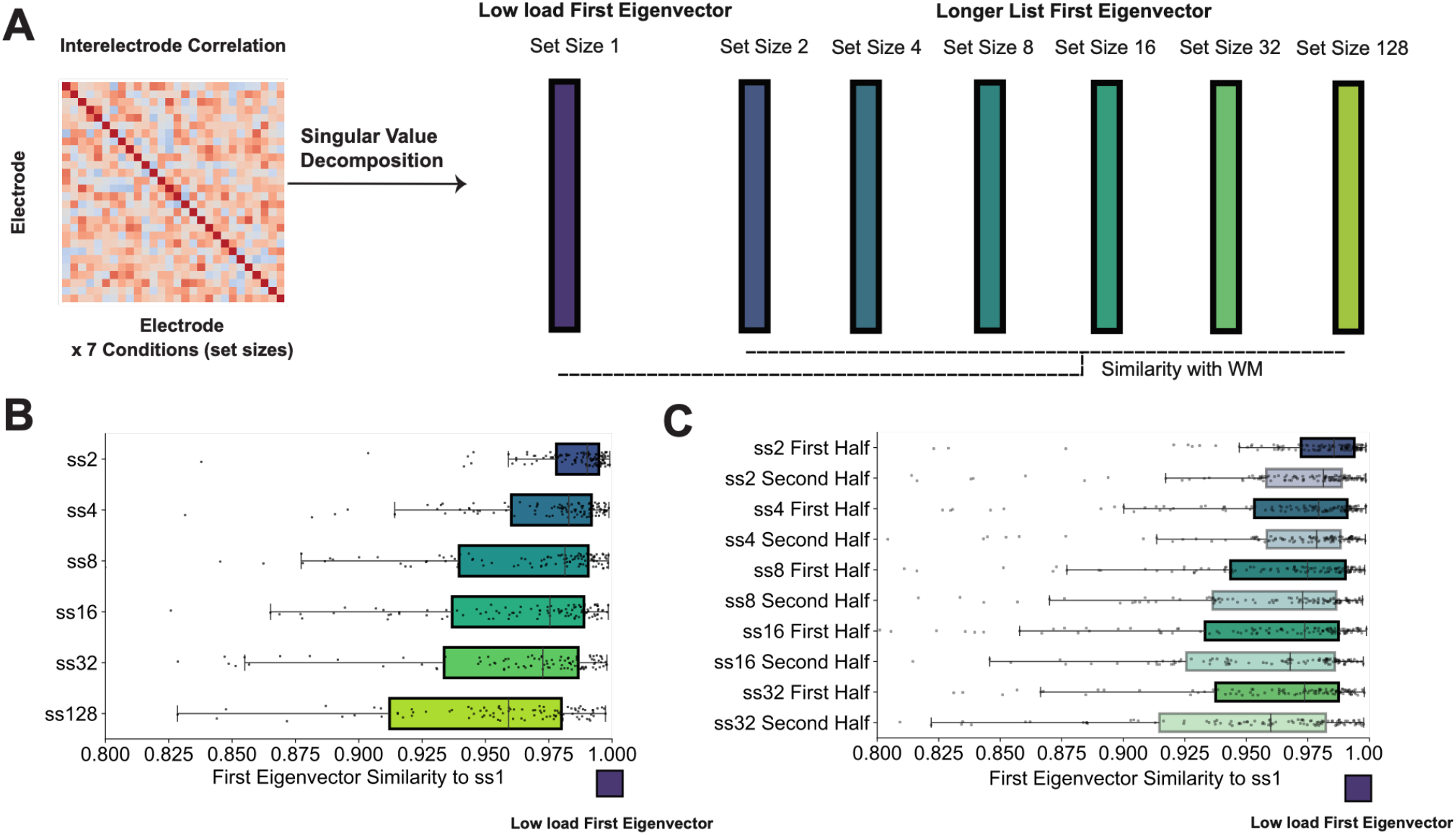
First eigenvector similarity on ICs underlie the shift from low to high load mode.

We used the first eigenvector derived from the low-load condition (set size 1) as a reference template and quantified its cosine similarity to the first eigenvectors obtained from each longer-list condition (**Fig. 3A**, dashed lines). As set size increased from 2 to 128 items, progressively recruiting high-load encoding processes, the similarity between the first eigenvector of each condition and the set size 1 low-load reference eigenvector decreased monotonically (**Fig. 3B**). Conditions requiring retention of small numbers of items (set size 2, set size 4) yielded first eigenvectors that remained highly similar to the low-load template (median pearson r > 0.97), while the largest set-size condition (set size 128) exhibited reduced similarity (median ∼0.925, *p* < 0.001) and greater inter-subject variability. The fact that the systematic shift from low-to-high load was captured by the first eigenvector suggests that the transition between memory systems is encoded along a single dominant axis of large-scale neural representation.

To examine whether the encoding state changed as mnemonic history accumulated while equating the amount of neural data contributing to each estimate, we again divided each condition into early and late portions, yielding 64 encoded items per estimate at every set size. We compared first- and second-half eigenvectors separately for each set-size condition **(Fig. 3C**). Within each condition, the two halves yielded highly concordant eigenvectors, indicating robust within-condition neural structure. Critically, the progressive decline in low-load template similarity was replicated across both session halves for all set sizes, confirming that the observed shift in dominant coordination structure reflects a genuine property of memory encoding rather than temporal drift or signal non-stationarity. Taken together, these findings suggest that the neural transition from low-load to high-load encoding reflects a trajectory in a low-dimensional state space.

### Rotation Angle Between Neural Subspaces Predicts Individual Memory Performance

Having established that the low-to-high load transition is captured by a low-dimensional reorganization of interelectrode correlation patterns, we next quantified the degree of representational change between conditions. Although mathematically equivalent to cosine similarity for normalized eigenvectors, angular distance provides an intuitive measure of how far the dominant coordination axis rotates away from the low-load reference. We therefore characterized the representational geometry more precisely using the rotation angle between IC patterns derived from each set-size condition and the low load reference (set size 1). As illustrated schematically in **Fig. 4A**, when two subspaces are highly similar, their rotation angle is small (e.g., ∼10°), reflecting a near-parallel alignment of dominant coordination axes; when representations are dissimilar, the rotation angle is large (e.g., ∼90°), indicating that the two subspaces are nearly orthogonal and thus capture fundamentally distinct neural coordination patterns.

**Figure 4.**
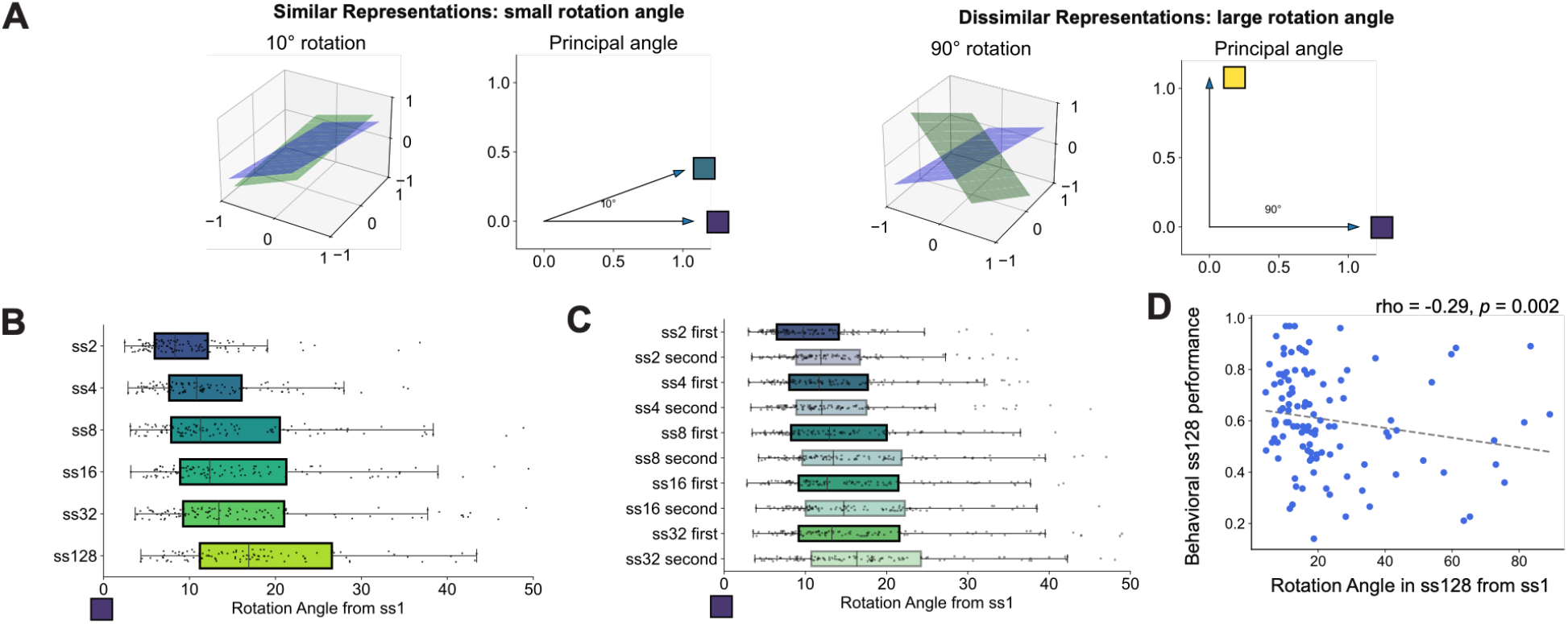
Rotation Angle drives the shift from low-load to high-load mode.

Applying this approach across all set-size conditions, rotation angles from the low load reference (set size 1) increased monotonically with list length (**Fig. 4B**). Suggesting that the effect was not driven by localized changes at specific electrodes, no individual electrode carried a significant weight in the transformation (ps > 0.28). These findings indicate that rotation of a neural subspace, rather than changes in any single electrode, is a key feature of the **adaptive reconfiguration** accompanying increasing mnemonic demands.

To examine whether the encoding state changed with accumulating mnemonic history while equating the amount of neural data contributing to each estimate, we again divided each condition into early and late portions as in our eaerlier analyses (**Fig. 4C**). Within every set-size condition, first- and second-half rotation angles were highly concordant, demonstrating that the neural subspace geometry is stable across the course of the experimental session. Critically, the monotonic increase in rotation angle with set size was consistently reproduced across both halves, ruling out explanations based on fatigue or other temporal confounds.

Although our prior analyses were focused primarily on within-subject effects, in the set size 128 condition, rotation angle also captured across-subject variability in behavioral performance. Participants who exhibited smaller rotation angles from the low-load subspace in set size 128 performed better on the subsequent recognition memory task (Spearman’s ρ=−0.29, p=0.002; **Fig. 4D**). These indviduals, who maintained a more low-load-like neural subspace even at set size 128, may have been better able to leverage existing coordination structure for high-load encoding, benefitting memory performance.

Taken together, these results establish that the low-to-high load encoding transition unfolds along a continuous rotational trajectory whose magnitude scales with memory load, and accounts for individual differences in LTM performance. This suggests that neural subspace rotation reflects a functionally significant reorganization of large-scale neural coordination associated with the efficiency of long-term memory encoding.

### Arousal Does Not Explain Differences in Subspace Rotations Across Subjects

A concern when interpreting condition-dependent changes in interelectrode correlation patterns is that they may be epiphenomenal consequences of incidental differences in arousal. For example, perhaps participants with large rotation angles between low-load and high-load modes showed declining arousal at longer lists. To test if arousal drove individual differences in neural rotation angle, we examined whether arousal, as indexed by pupil size, predicted rotation angles in the high-load condition. Pupil responses were separated into tonic and phasic components to capture sustained and transient changes in pupil size, respectively. Tonic pupil size was quantified as the baseline pupil diameter from −200 ms to 0ms encoding image onset, whereas phasic pupil change was calculated as the image-evoked change (300-800 ms) in pupil size relative to this −200 to 0 ms baseline. Across 73 participants that had quality eye tracking data, neither tonic nor phasic pupil responses were associated with individual differences in neural rotation (tonic: Spearman’s ρ = 0.003, p = 0.982; phasic: ρ = −0.038, p = 0.749) or LTM performance (tonic: Spearman’s ρ = −0.135, p = 0.256; phasic: ρ = −0.001, p = 0.988). Thus, although pupil responses may reflect changes in encoding states within individuals, they did not explain individual differences in memory performance as rotation angles did.

### EEG Signal Amplitude Does Not Explain Subspace Rotations

We finally investigated whether individual differences in subspace rotation angle could be explained by changes in overall EEG signal amplitude distributions. We examined Current Source Density (CSD) scalp topographies, which characterize the spatial profile of oscillatory power across the scalp, separately for each set-size condition (**Fig. 5**). CSD topographies were highly similar across all conditions (*p*s > 0.16). The characteristic frontal-to-occipital gradient was preserved from set sizes 1 through 128, with no systematic reorganization of amplitude distributions. It is therefore unlikely that the IC eigenvector rotations were simply consequences of cross-condition differences in electrode amplitude landscapes.

**Figure 5.**
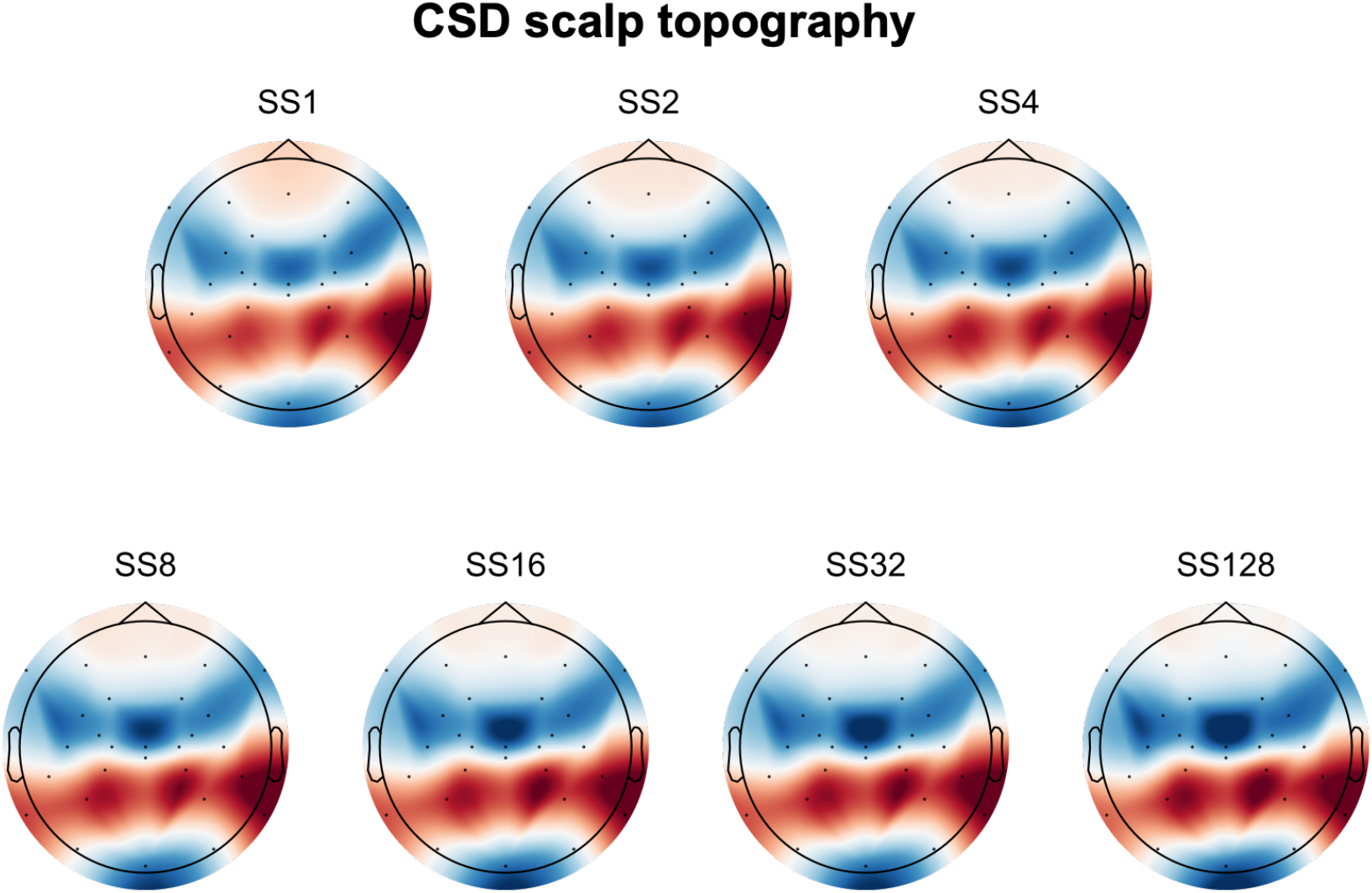
Scalp Topography did not drive the change from low-load to high-load encoding.

## Discussion

The present findings provide converging evidence for adaptive reconfiguration of the brain state associated with memory encoding as mnemonic demands increase. Rather than simply repeating the same encoding architecture as lists become longer, neural representations progressively shifted away from the configuration observed at low loads and toward a distinct high-load configuration. This conclusion is supported by three complementary observations: (1) interelectrode correlation (IC) patterns progressively shifted with set size, (2) this shift was captured by a low-dimensional change in neural coordination, and (3) the magnitude of neural subspace rotation was related to individual differences in LTM performance. Together, these results favor the adaptive reconfiguration account over repeated loading.

At small set sizes, neural representations closely resembled the low-load template. Specifically, set sizes 2 and 4 showed significantly greater similarity to the set size 1 template, indicating that encoding remained within a common neural configuration at or near canonical WM capacity ^1,3^. Importantly, the split-half analyses showed that this organization was maintained throughout the set size 4 lists, rather than being restricted to their earliest items. Thus, within capacity, encoding remained relatively stable across successive items.

A different pattern emerged as mnemonic demands accumulated beyond capacity. At set size 8, the first four items remained more similar to the low-load template, whereas the latter four showed an intermediate pattern. At set size 16, the later items shifted further toward the high-load template, and by set size 32, later-list encoding was clearly more similar to the high-load configuration. Thus, increasing demands did not simply produce repeated instances of the same low-load encoding pattern. Instead, neural organization progressively changed as items accumulated, providing direct evidence for adaptive reconfiguration. The within-list results further suggest that this reconfiguration unfolds as mnemonic demands build, rather than being determined solely by the nominal set size.

This pattern is difficult to reconcile with a strong repeated loading account. If long lists were encoded by repeatedly cycling through the same encoding architecture used for shorter lists, a 16-item list, for example, should resemble repeated applications of the configuration observed for a 4-item list. IC patterns should therefore remain relatively stable across list lengths, aside from changes in the amount of loading. Instead, neural patterns systematically diverged from the low-load configuration as lists lengthened. The intermediate patterns at set sizes 8 and 16 suggest that this reconfiguration is graded rather than an abrupt switch between two independent systems. Thus, increasing mnemonic demands appear to alter the organization of encoding itself, rather than simply increasing the amount of processing within a fixed architecture.

This reconfiguration was also remarkably low-dimensional. Across memory loads, SVD revealed that changes in IC structure were largely captured by the first eigenvector. The progressive decrease in similarity between the first eigenvector for each condition and the low-load reference indicates that the shift in neural organization was concentrated along a dominant dimension rather than distributed across many independent dimensions. Rather than requiring a wholesale restructuring of the neural state space, adaptive reconfiguration may therefore involve movement along a constrained trajectory through neural state space as mnemonic demands increase. The global low-dimensional geometries we observed during visual memory encoding raise the question of whether such dimensionality persists across scales and species ^14^. Future work should examine whether local geometric dynamics drive this global shift during visual memory encoding.

The rotation-angle analysis provides a complementary measure of this trajectory. Neural patterns rotated progressively farther from the low-load reference as set size increased, consistent with a graded reorganization rather than a categorical switch. Importantly, this neural reconfiguration was functionally relevant: at set size 128, larger rotations were associated with poorer LTM performance, whereas more low-load-like neural geometry predicted better performance. Successful long-list encoding may therefore depend not simply on maximizing reconfiguration, but on achieving an effective balance between the original encoding architecture and its adaptation to increased mnemonic demands.

Together, these findings extend capacity-based accounts of memory by showing that the limit of WM is accompanied by a systematic transformation in large-scale neural coordination. Long lists are not simply encoded as repeated applications of the same WM architecture. Instead, as mnemonic demands accumulate, the brain progressively reconfigures its encoding dynamics along a low-dimensional trajectory. Adaptive reconfiguration therefore provides a framework for understanding how a common memory system can flexibly alter its organization when the demands of encoding exceed the capacity of its initial operating regime.

## Method

### Participants

One hundred and eleven participants took part in the study (mean age = 25.1 years). Sample size was determined using G*Power, aiming to achieve 80% power to detect a small effect size (r = 0.3) at the conventional alpha level of .05. All participants reported normal or corrected-to-normal vision and normal color perception, and no history of neurological disorder. Study procedures were approved by the relevant University of Chicago Institutional Review Board and participants were compensated for their participation.

### Stimuli and Procedure

Images were selected from the THINGS dataset ^15^, a database with 1854 unique concepts and 26107 images. One exemplar was selected from each concept so that no repeated concept was used throughout each session (i.e., apple versus avocado). The recognition memory procedure for all set sizes (1, 2, 4, 8, 16, 32 and 128) followed established protocols in the literature and consisted of separate study and test phases ^16,17^. The experiment included set sizes of 1, 2, 4, 8, 16, 32, and 128. Each encoding condition consisted of a total of 128 trials. This was achieved by varying the number of blocks across set sizes, such that participants completed 128 blocks in the set size 1 condition, with progressively fewer blocks as set size increased, down to a single block in the set size 128 condition (see **Fig. 1**).

In the set size 1 condition, participants viewed a single image presented at the center of the screen for 800 ms during the encoding phase. Stimuli were displayed against a gray background (90.0 cd/m^2^), and a white fixation cross was superimposed on the image to encourage central fixation and discourage eye movements. The interstimulus interval was randomly jittered between 250 and 400 ms.

Following the encoding phase, participants completed a recognition test trial in which a single image was displayed at the center of the screen for 800 ms with a central white fixation cross. To minimize motor and ocular artifacts, participants were instructed to remain still while the fixation cross was visible. After 800 ms, the fixation cross disappeared and the image remained on screen until a response was made. Half of the test images had been presented during the encoding phase (old), and the remaining half were novel (new). The order of old and new images was randomized across trials. Participants indicated whether each image was “old” or “new” using designated keyboard keys (“z” for old; “/” for new).

All other set size conditions followed the same general structure, differing only in the number of images presented during the encoding phase and the corresponding number of test trials. Block order was randomized across participants.

### Preprocessing and artifact rejection of EEG signals

Participants were positioned in an electrically isolated booth, with their heads stabilized using a cushioned chinrest placed 74 cm from the display screen. Electroencephalographic (EEG) signals were recorded via 30 active silver/silver chloride (Ag/AgCl) electrodes integrated into a stretchable cap (actiCHamp system, Brain Products, Munich, Germany), arranged according to the international 10-20 placement standard (electrode sites: Fp1, Fp2, F7, F8, F3, F4, Fz, FC5, FC6, FC1, FC2, C3, C4, Cz, CP5, CP6, CP1, CP2, P7, P8, P3, P4, Pz, PO7, PO8, PO3, PO4, O1, O2, Oz). Additional electrodes were adhered to the left and right mastoids using adhesive stickers, and a ground electrode was integrated at the Fpz site within the cap. EEG signals were initially referenced to the right mastoid and subsequently re-referenced offline to the average of both mastoids. The signal was bandpass filtered (0.01–80 Hz) with a 12 dB/octave roll-off and digitized at a sampling rate of 500 Hz. All electrode impedances were maintained below 10 kΩ.

To track ocular activity such as blinks and saccades, both electrooculographic (EOG) and eye-tracking data were recorded. EOG was captured using five passive Ag/AgCl electrodes: two for vertical EOG (above and below the right eye), two for horizontal EOG (approximately 1 cm lateral to each eye), and one ground electrode on the left cheek. Eye position was also monitored with a desktop-mounted EyeLink 1000 Plus system (SR Research, Ontario, Canada), operating at a sampling rate of 1,000 Hz.

To enhance spatial specificity and reduce volume conduction effects, we applied the surface Laplacian (current source density, CSD) transformation to the EEG data using the spherical spline method. This approach estimates the second spatial derivative of the scalp potential, effectively acting as a spatial high-pass filter and improving signal-to-noise ratio (SNR) for inter-electrode correlation analyses. CSD was computed using the compute_current_source_density function from MNE-Python ^18–20^ based on the 3D locations of 28 scalp electrodes.

### Artifact Rejection

For horizontal eye movements, we employed a sliding-window algorithm to identify horizontal eye movements using both horizontal electrooculogram (HEOG) signals and eye-tracking gaze data. For the HEOG-based detection, a split-half sliding-window method was applied with a 100 ms window moving in 10 ms increments. An eye movement was flagged if the voltage difference between the two halves of the window exceeded 20 µV. This HEOG-based artifact detection was used only in trials where eye-tracking data were unreliable or unavailable. In parallel, eye-tracking rejection was carried out by analyzing horizontal (x-axis) and vertical (y-axis) gaze positions using the same 100 ms window and 10 ms step size. A trial was excluded if eye position shifted more than 0.5° of visual angle within any given window.

To detect blinks, we applied a sliding-window analysis to the vertical EOG signal using an 80 ms window and a 10 ms step size. A blink was marked when the voltage change across the window exceeded 30 µV. Complementarily, we identified blink periods in the eye-tracking data by locating segments where positional data were missing, indicating that the eyes were closed. For set size 32 trials, the overall rejection rate is 0.78% (1.0 trials). For set size 128 trials, the overall rejection rate is 2.24% (2.9 trials).

### Inter-electrode EEG Correlations (IC)

We measure correlations between the mean trial time courses between all pairs of electrodes ^12^. To do so, we averaged the raw amplitude at each of our non-reference 28 electrodes across all trials from time points 0 (the onset of the memory array) to 800 ms, which is the longest artifact-free time for each trial during the coding phase. Since we have four set size 32 blocks and one set size 128 block, the number of trials during the encoding phase was the same for both conditions, so the signal-noise-ratio of the resulting event-related potential (ERP) was similar between our two different conditions. We next computed the Pearson correlation of this trial-averaged ERP for all pairwise electrodes for each participant separately. For each participant, this resulted in a 28 x 28 matrix of the correlation between the time course of each electrode to each other electrode.

### Singular value decomposition (SVD) of IC matrices

To determine whether condition differences in IC structure were high-dimensional or low-dimensional, we decomposed each participant’s IC matrix using singular value decomposition. For each set-size-specific correlation matrix *X*, we computed

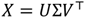

where the columns of *U* and *V* are orthogonal singular vectors and Σ contains the singular values in descending order. Because the IC matrices are symmetric, the left and right singular vectors are equivalent to the eigenvectors of the matrix. We focused primarily on the first eigenvector (that is, the singular vector associated with the largest singular value), as this component captures the dominant axis of coordinated variance across electrodes. To test whether the WM-to-LTM transition was captured by this dominant dimension, we used the first eigenvector from the set size 1 condition as the WM reference and compared it with the first eigenvector from each larger set-size condition.

### Eigenvector similarity analysis

Similarity between first eigenvectors was quantified using cosine similarity. For two unit-length For two unit-length eigenvectors *u* and *v*, similarity was computed as

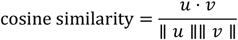

with values closer to 1 indicating highly similar dominant neural structure and lower values indicating greater divergence. Because the sign of eigenvectors is arbitrary, vectors were sign-aligned before comparison so that similarity reflected shared geometry rather than sign inversion. This analysis allowed us to test whether the dominant coordination axis became progressively less similar to the WM template as set size increased.

### Rotation angle between neural subspaces

To provide a geometrically interpretable measure of representational change, we quantified the rotation angle between the dominant neural subspace of each condition and that of the WM reference condition (set size 1). Using the cosine similarity between first eigenvectors, the rotation angle *θ* was defined as

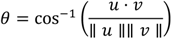

where *u* is the first eigenvector from the set size 1 IC matrix and *v* is the first eigenvector from the comparison condition. Smaller angles indicate close alignment between neural subspaces, whereas larger angles indicate greater representational rotation away from the WM-dominant encoding geometry. We computed this angle separately for each participant and set-size condition, enabling us to test whether neural subspace rotation scaled monotonically with memory load and whether it predicted behavioral performance.

### Current source density (CSD) topography

To assess whether the dominant IC structure could be trivially explained by condition differences in scalp amplitude distribution, we computed current source density (CSD) scalp maps for each set-size condition. CSD was estimated from the scalp voltage data using the surface Laplacian, which computes the second spatial derivative of the voltage field across neighboring electrodes. This transformation attenuates volume conduction and reference-dependent effects while sharpening the local spatial distribution of cortical activity. In practice, for each participant and condition, trial-averaged scalp voltages during the encoding period were transformed into reference-free CSD values using a standard spherical spline Laplacian procedure. The resulting maps indexed the spatial topography of local current flow across the scalp. We then visually and quantitatively compared CSD topographies across set-size conditions to determine whether the IC-based subspace rotations could be attributed to systematic differences in scalp amplitude layout rather than genuine changes in inter-electrode coordination geometry.

## Ethics

Study procedures were approved by the relevant University of Chicago Institutional Review Board.

## Data and Code Availability

The repository will be made available upon paper acceptance on Open Science Framework.

## Acknowledgement

This research was supported by funding from the National Institute of Mental Health (Grant ROIMH087214 awarded to Edward K. Vogel), and the Office of Naval Research (Grant N00014-12-1-0972 awarded to Edward K. Vogel; Grant MURI N00014-23-1-2768 to Monica D. Rosenberg and Edward K. Vogel).

## Author Contributions

Chong Zhao played a lead role in conceptualization, data curation, formal analysis, investigation, methodology, writing–original draft, and writing– review and editing. Edward K. Vogel played a lead role in conceptualization, funding acquisition, resources, supervision, and writing–review and editing. Monica D. Rosenberg played a lead role in conceptualization, formal analysis, investigation, methodology, funding acquisition, resources, supervision, and writing–review and editing.

## AI Acknowledgement Statement

During the preparation of this work, the authors used ChatGPT and Claude to assist with manuscript revision and ChatGPT to streamline the color schemes and formatting of the figures. The authors carefully reviewed and revised all materials and take full responsibility for the final content.

